# Evidence that loss of consciousness under anesthesia is not associated with impaired stimulus representation in auditory cortex

**DOI:** 10.1101/213355

**Authors:** Matthew I. Banks, Bryan M. Krause, Nicholas S. Moran, Sean M. Grady, Jeremiah Kakes, Daniel J. Uhlrich, Karen Manning

## Abstract

The mechanism whereby anesthetics cause loss of consciousness (LOC) is poorly understood. Current theories suggest that impaired representation of information in cortico-thalamic networks contributes to LOC under anesthesia. We sought to determine whether such changes are present in auditory cortex using information theoretic analysis of multiunit responses in rats. We tested the effects of three agents with different molecular targets: isoflurane, which acts at multiple pre- and postsynaptic loci, propofol, which acts primarily on GABA_A_ receptors, and dexmedetomidine, an α_2_ adrenergic agonist. We reasoned that changes in the representation of sensory stimuli causative for LOC would be present regardless of the molecular target of the anesthetic. All three agents caused LOC, as assayed by the loss of righting reflex (LORR). We presented acoustic stimuli that varied across a wide range of temporal and spectral dynamics under control, sub-hypnotic (i.e. dose too low to cause LORR), just-hypnotic (a dose just sufficient to cause LORR) and recovery conditions. Changes in mutual information (MI) between the stimulus and spike responses under anesthesia diverged in two ways from predictions of a model in which stimulus representation is impaired upon LOC. First, the sign of changes in MI was agent-specific: MI increased under dexmedetomidine, while it decreased under isoflurane and propofol. Second, there was no consistent change in MI when transitioning from sub-hyptnotic to just-hypnotic doses: for none of the agents did MI decrease at the higher dose, and in some cases MI actually increased relative to the sub-hypnotic dose. Changes in MI under anesthesia were strongly correlated with changes in precision and reliability of spike timing, consistent with the importance of temporal stimulus features in driving auditory cortical activity. These data indicate that primary sensory cortex is not the locus for changes in information representation causative for LOC under anesthesia.

## Introduction

What changes in the brain upon LOC under anesthesia? This question has broad implications both for improving clinical practice as well as for understanding the neural basis of consciousness. Theoretical considerations suggest that anesthetics impact cortical representations of information and its communication across brain regions (Alkire et al., 2008, Tononi, 2004 #6447, Mashour, 2013 #7652). Several studies using EEG recordings and functional magnetic resonance imaging have provided indirect evidence for these models in patients and volunteers (Imas et al., 2005;Peltier et al., 2005;Alkire, 2008;Lee et al., 2009;Ku et al., 2011;Liu et al., 2011;Schrouff et al., 2011;Boly et al., 2012;Jordan et al., 2013;Lee et al., 2013a;Lee et al., 2013b;Blain-Moraes et al., 2014;Mashour, 2014). However, these studies focused on information and connectivity in higher order cortical areas in the absence of sensory stimulation, and thus the representation of sensory information in primary sensory areas has not been explored in detail.

Indeed, primary sensory cortex may be excluded from the cortico-thalamic network mediating consciousness in some models (Logothetis et al., 1996;Tononi, 2004;Watanabe et al., 2011), and there are suggestions that during anesthesia LOC, sensory information from the periphery activates primary sensory cortex (Guo et al., 2012;Durand et al., 2016), but fails to engage higher order cortical processing (Liu et al., 2011). We recently observed that synaptic responses to auditory stimuli and stimulation of auditory thalamocortical afferents were maintained in A1 under isoflurane (Raz et al., 2014), suggesting that auditory stimulus representation at the single cell level in auditory cortex should be relatively unaffected.

In contrast to the straightforward predictions of the studies cited above, there is a large body of literature suggesting that sensory responses in A1 are affected dramatically by anesthetic agents. For example, imaging and EEG studies in both experimental animals and humans show that sensory responses in cortex are suppressed under anesthesia, even at sub-hypnotic doses, (Schwender et al., 1993;Dueck et al., 2005;Plourde et al., 2006), and we observed suppression of thalamocortical synaptic responses in brain slices, albeit less than observed for corticocortical synaptic responses (Raz et al., 2014). Anesthetics have been reported to suppress activity in thalamus both *in vivo* (Alkire et al., 2000) and in brain slice preparations (Ries and Puil, 1999), observations that are consistent with the thalamic switch hypothesis (Alkire et al., 2000) in which anesthetics cause LOC by disconnecting the cortex from the periphery. Thus, we still lack a basic understanding how anesthetics affect encoding and processing of sensory responses, a critical step in elucidating the mechanisms of loss and recovery of consciousness under anesthesia (Hudetz et al., 2015).

We investigated the cortical effects of three anesthetics that can each independently produce LOC: propofol, dexmedetomidine and isoflurane. The receptor targets of propofol are GABA_A_ receptors, whereas dexmedetomidine targets α_2_ adrenergic receptors, and isoflurane has a number of molecular targets including GABA_A_ and NMDA receptors and two-pore-domain K^+^ channels (Alkire et al., 2008, Franks, 2008 #7064, Rudolph, 2004 #5754). Both isoflurane and dexmedetomidine have presynaptic effects on glutamate release (MacIver et al., 1996, Chiu, 2011 #7422). In spite of the diversity of their chemical structure and molecular targets, isoflurane, propofol and dexmedetomidine all produce LOC and all have direct actions in neocortex (Alkire et al., 2000, Boly, 2011 #7243, Ferrarelli, 2010 #7209, Hentschke, 2005 #7263, Murphy, 2011 #7265, Ries, 1999 #5156, Velly, 2007 #7264). We reasoned that changes in sensory representation that are causative for anesthesia LOC would be observed at just-hypnotic doses of all three agents.

We tested this hypothesis by recording unit activity in auditory cortex of chronically-implanted rats in response to acoustic stimuli under control, sub-hypnotic and just-hypnotic drug conditions. We computed standard rate and timing measures of unit activity, but specifically assayed the representation of information in spike trains using information-theoretic analyses (Nelken and Chechik, 2007, Schnupp, 2006 #6260, Shannon, 1948 #6446). If anesthetic suppression of information in cortical networks is tightly linked to LOC, these analyses should show dramatic differences between stimulus representation at sub-hypnotic versus just-hypnotic doses. We did not observe this expected relationship, suggesting that changes in stimulus representation in sensory neocortex are not causative for anesthesia LOC.

## Methods

All experimental protocols conformed to American Physiological Society/National Institutes of Health guidelines and were approved by the University of Wisconsin Animal Care and Use Committee. The data reported here were recorded from 6 female ACI rats (Harlan; 139 – 204 gms) housed in groups of 3-4 animals prior to surgery and individually afterward, and maintained in a humidity and temperature-controlled environment on a reverse 12:12 light dark cycle so that recordings performed during the day were during the animals’ ‘active’ period.

### Surgeries

Animals were implanted with multichannel microwire arrays and intrajugular catheters before commencing recording sessions. Both types of surgery were performed under aseptic conditions and isoflurane anesthesia (1.5 - 2% in 50% O_2_/50% room air). Rats were premedicated with Meloxicam (1 mg/kg SQ) to manage pain and swelling and Bupivicaine (1.0mg/kg SQ) was applied locally at the site of incision for pain. Rats were kept on an infrared heating pad throughout surgery and recovery to maintain core temperature at 37±0.5°C. Rats were hydrated with 0.9% saline via subcutaneous injections of 2 - 10 ml/kg and ophthalmic ointment used to prevent dehydration of the cornea. Animals were treated postoperatively for pain (buprenorphine 0.02 mg/kg SQ and meloxicam 1 mg/kg SQ) for three days following surgery and checked daily for indications of infection or any discomfort. Following surgery and throughout the recording period, animals were maintained on the same reversed light/dark schedule.

### Implantation of indwelling intrajugular catheter

Chronic implantation of intrajugular catheters was performed to allow for administration of intravenous anesthetics. Following isoflurane induction, 2 surgical sites, the right ventral neck and chest region and a dorsal area between the shoulders, were shaved and then scrubbed with 70% isopropyl alcohol and Betadine™ solution. Rats were placed dorsally on a heated pad. An 8–10-mm ventral paramedian skin incision was made superficial to the carotid artery and parallel to the trachea.

The right, external jugular vein was made accessible using blunt dissection of surrounding tissue and kept moist with sterile water to prevent collapse. Two ligatures were placed at least 2 mm apart, both distal to the origin of the axillary vein. After the rostral ligature was tightened, the right jugular was venotomized between the 2 ligatures with a 27-gauge needle. A silastic catheter filled with heparinized saline (20 mg/ml) and attached to a 1 cc syringe was inserted 7–10 mm into the jugular vein. After catheter placement, a small amount of blood was drawn into the catheter and then flushed with heparinized saline to ensure catheter patency. The catheter was secured and the vein occluded by tightening the proximal ligature and by tying additional knots above and below the silastic collar. At this point, the catheter was disconnected from the 1-cc syringe. After exiting the jugular vein, the distal end of the catheter was directed subcutaneously along the neck and externalized through a small dorsal skin incision between the scapulae. Care was taken during this phase of the surgery to ensure that the catheter did not become twisted. The incision was small enough that the skin stretched tightly around the catheter. The ventral incision was then closed with a tissue adhesive (Vetbond™, No. 1469, 3M Animal Care Products) or 6-0 proline and the dorsal incision sealed to the slip-luer syringe tip (i.e., No. 5). Immediately after surgery, rats were infused with heparinized saline (20 mg/ml) for 30 min (0.0034 ml/min) to replenish bodily fluids.

### Implantation of electrode arrays

Following isoflurane induction, the skin over the head was shaved and disinfected with Betadyne and the skull exposed via midline skin incision. The skull was cleared of connective tissue and dried with ethyl alcohol. Skull screws were placed bilaterally into the frontal plates and parietal plates, and ipsilaterally in the occipital plate, the latter of which served as the recording ground. A craniotomy was then made to allow access to auditory cortex. To record unit activity, we implanted 2×8 (rows x columns) tungsten microwire arrays (polyimide insulated, wire diameter 33 µm, impedance 1 MΩ, column spacing 250 µm, row separation 500 µm; Tucker Davis Technologies, Alachua, FL), with the long axis of the array oriented rostro-caudally (Supplementary Figure 1A). The cortical target was A1, though in some animals the caudal-most 1 – 2 electrodes in each row were determined to be in PAF, and in one animal the electrodes were entirely located in AAF. Structures were targeted stereotaxically (Doron et al., 2002;Paxinos and Watson, 2007;Polley et al., 2007) and post-hoc histological analysis (see below) was used to verify electrode location. A 1.85 mm x 1 mm craniotomy (R-C x M-L) on the left side, centered at 5.25 mm caudal to Bregma, 5.9 mm off the midline was made. The electrode was implanted using an angled dorsal approach, with the angle off the midline = 20°. After reflecting the dura, the electrode array was inserted using a remotely controlled hydraulic microdrive (**Model, Company, City, State**) until the recording sites were located in the appropriate region (2.4 – 3.25 mm), as judged by responses to acoustic stimuli recorded during the implantation process. The craniotomy was filled with silicone elastomer (Kwik-Sil, World Precision Instruments, Sarasota, FL) and the array and headstage connector were cemented to the skull and skull screws using dental acrylic **(Model, Company, City, State)**. Animals were allowed to recover 5-7 days before the first recording session.

**Figure 1.**
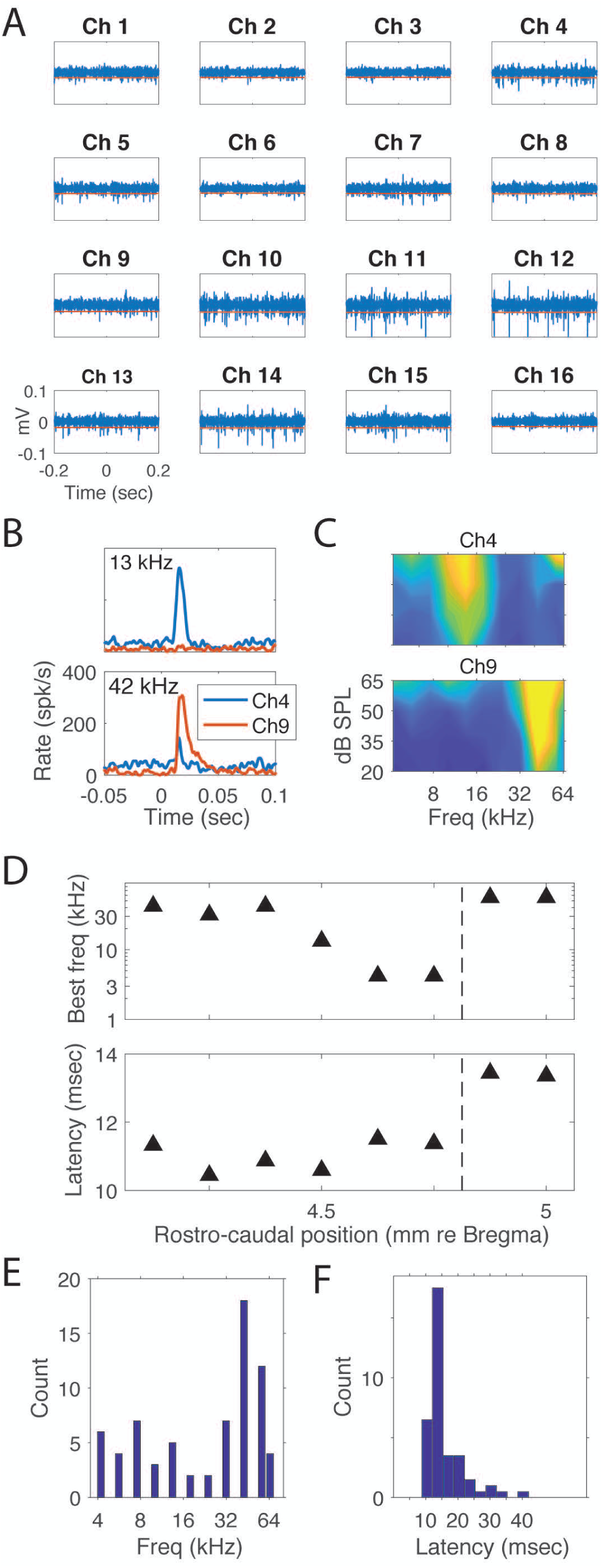
Example of unit activity and frequency tuning for single electrode and an entire array. A) 16 channels of single trial raw data with threshold lines. B) Example histograms for two electrodes. C) FRAs for two electrodes. D) Example of tuning (top) and latency (bottom) as function of R-C location in one animal. E) Histogram of BFs for all electrodes. F) Histogram of pure tone spike latencies at BF for all electrodes.

### Electrophysiological recordings

Experiments on each animal were performed over a period of three weeks, 3 – 4 recording sessions per week. For each week, one of three stimulus sets was selected (click trains, vocalizations, FM sweeps; see below) and used for all recordings during that week. Each day a different drug was selected (isoflurane, dexmedetomidine, propofol; see below), with 48 hours between recording sessions. All recordings took place inside an anechoic sound-attenuation chamber (Industrial Acoustics Company, Inc., Bronx, NY). On the day of the experiment, animals were placed inside the chamber in a gas-tight acrylic enclosure (20 × 19 × 11 cm) and allowed to accommodate to the enclosure for one hour. The enclosure had gas inflow/outflow and sampling ports used to provide room air in all experiments and to deliver isoflurane during that subset of the experiments (see below). Animals were kept warm using a heating pad placed in the bottom of the enclosure. Microwire signals were access by connecting a 16 channel headstage (**Model**, Tucker Davis Technologies, Alachua, FL) with a flexible tether to a 16 channel connector on the animal’s head. The headstage and tether did not impair the animal's ability to move about the enclosure, though the small size of the enclosure ensured that the animals were in approximately the same position relative to the overhead speaker. Responses were bandpass-filtered at 1-7500 Hz, digitized at 24kHz and amplified at 5000 - 20000X (RZ5, Tucker Davis Technologies).

Free-field acoustic stimuli were presented via a small speaker (TDT-ES1, Tucker Davis Technologies) mounted in the top (‘ceiling’) of the enclosure, pointed downwards. A microphone (#4016, ACO Pacific, Inc., Belmont, CA) placed approximately 4 cm from the speaker was used for calibration, and stimuli presented at approximately 20 - 70 dB SPL assuming the animal’s head was this distance from the speaker. Since the animal was unrestrained, actual stimulus levels on each trial varied slightly. Speaker output varied by <±10 dB SPL over the range 4 – 60 kHz. During the first recording session, frequency response areas were obtained using pure tone stimuli (50 msec duration, 5 msec rise/fall, 4 kHz – 60 kHz in 11 log-spaced steps, 20 – 65 dB SPL). During subsequent recording sessions, stimuli consisted of click trains (0.1 ms clicks, 20 dB attenuation, 1 second train, interclick intervals 10 – 333 msec), upward and downward frequency modulated (FM) sweeps (250 ms duration, range 4 kHz – 32 kHz, f_Low_ = 4 kHz – 32 kHz, f_High_ = 8 kHz – 64 kHz) and a set of twelve animal vocalizations recorded from birds, insects, and rodents (Avisoft Bioacoustics; www.avisoft.com) that have distinct spectro-temporal properties (Suplementary Figure 2). Specific FM sweep frequencies were chosen for each animal to span the frequency response areas of the recording sites. Vocalizations were either 250 msec or 500 msec and six stimuli were selected for each animal from the set of twelve to overlap the frequency response areas of the recording sites. Stimuli were generated and recordings obtained using commercial software (Brainware, RPVDX, Tucker-Davis Technologies). Spontaneous activity was also recorded for five minutes during each treatment condition.

**Figure 2.**
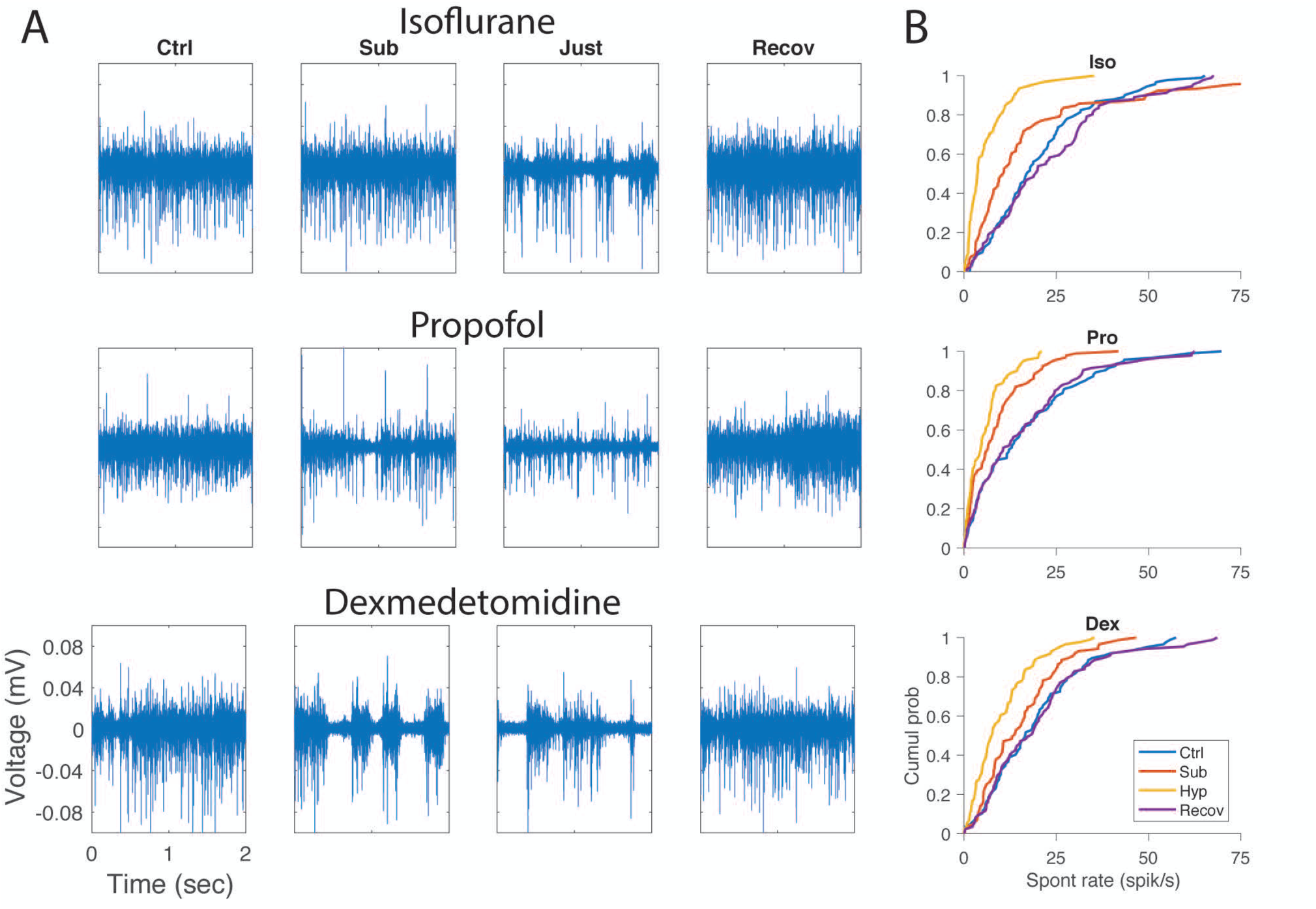
Effects of anesthetics on spontaneous rate. A) Raw MUA signal from the same channel in one animal under control (left column), subhypnotic (second column), just-hypnotic (third column) and recovery (right column) for isoflurane (top row), propofol (middle row) and dexmedetomidine (bottom row). B) Cumulative probability distributions of spontaneous firing rate across all clusters and all animals.

On each recording day, two sets of control data were recorded, consisting of 150 – 200 trials of 6 – 7 stimuli from the selected stimulus type recorded over ~20 minutes. The two stimulus-evoked datasets were compared by eye and if noticeably different in responsiveness or firing rate, a third control data set was collected. The anesthetic agent selected for that day was then applied as described below. Finally, three sets of recovery data were then collected over ~1 hour following cessation of the anesthetic agent. For all analyses presented below, we are comparing data collected during the final control period, the sub-hypnotic and hypnotic recording periods, and the final recovery period.

### Anesthetic administration

Dexmedetomidine and propofol were administered intravenously during electrophysiological recordings using a syringe pump (**Model, Company, City, State**) and flexible lightweight tubing that did not impede the animal’s mobility. Isoflurane was applied in the gas phase in room air using a calibrated vaporizer and the concentration measured using a gas monitor (Multigas Monitor 602, Criticare Systems, Waukesha, WI). In each session, we recorded responses during four treatment conditions: awake (~1 hour), followed by two drug periods (sub-hypnotic and then just-hypnotic doses; ~40 minutes each) and recovery (~90 minutes). Each drug period consisted of a 20 min equilibration period followed by a 20 minute steady-state period. Responses were recorded throughout, but only data obtained during the steady-state period are presented here. During the equilibration period for the just-hypnotic dose, animals were tested every five minutes for LOC by tilting the anesthesia chamber by 30 degrees to detect loss of righting reflex (LORR), and the drug dose adjusted until LORR occurred. Although righting reflex is only a surrogate for awareness in rodents, there is strong correlation in dose dependence between loss of righting reflex in animals and LOC in humans across a remarkably diverse array of anesthetic agents (Franks, 2008). Drug doses were targeted to be subhypnotic and ‘just-hypnotic’ (MAC_Awake_ or Cp_Awake_)(Bol et al., 1999;Tung et al., 2008). Doses were as follows: isoflurane: subhypnotic 0.4% in room air, just-hypnotic: 0.8-1.0%, as needed to achieve LORR; propofol: subhypnotic 450 µg/kg/min, just hypnotic 600 – 700 µg/kg/min, as needed to achieve LORR; dexmedetomidine: subhypnotic 1.33 µg/kg/min for 10 minutes followed by 0.27 µg/kg/min, just-hypnotic 2.00 µg/kg/min for 10 minutes followed by 0.4 – 0.5 µg/kg/min, as needed to achieve LORR. For the recovery period, isoflurane and propofol were turned off and animals regained consciousness within about 5 minutes. For dexmedetomidine, we found that recovery could take more than an hour after turning off the drug, so we used a reversal agent [Antisedan (Atipamezole), 1 mg/kg, IP] to speed this process. Animals regained consciousness within 5 minutes after injection of the reversal agent.

### Data analysis

Raw data were filtered 500 – 3000 Hz to isolate unit activity. To eliminate noise common to all channels, we subtracted from each data channel the mean of the other channels as a common reference subtraction procedure. Because spikes were detected as negative excursions of the data, positive values of the mean were set to zero to eliminate introducing spurious signals recorded on individual channels to the rest of the channels. Spikes were detected as negative-going threshold crossings, where the threshold was set to 5/0.6745 times the median of the baseline noise (Quiroga et al., 2004). Thresholds were set independently for each channel. However, because of reduced motor activity in the drug conditions and due to the suppressive effect of anesthetics on spontaneous spiking activity, the noise levels during the drug conditions were substantially lower than in control or recovery. To assay changes in activity across all treatment conditions, the threshold was set based on data from the control condition, and applied to both drug conditions and the recovery condition.

We assayed effects of anesthetics on unit activity by measuring spontaneous and driven firing rates and first spike latencies. Spontaneous rate was calculated from the pre-stimulus window (usually 500 msec), and driven rate was calculated as the difference in firing rate during the stimulus minus the spontaneous rate. First spike latency was estimated as the first time point after the stimulus at which the spike pdf exceeded its pre-stimulus mean +2SD. To measure spike timing precision across trials in a single spike cluster for the vocalization stimuli, which were not periodic, we adapted the spike time tiling coefficient (STTC) (Cutts and Eglen, 2014), which is less dependent on firing rate than the correlation index (Joris et al., 2006) or other related measures. To evaluate spike timing during click trains, we used the STTC as well as the vector strength (VS) (Goldberg and Brown, 1969) and evaluated its significance via the Rayleigh distribution. For STTC and VS calculated from click train data, we ignored the first 10% (i.e. 100 msec) of the response period to avoid onset effects, and we included only spikes occurring between 5 and 40 msec after each click in the train.

We used an information theoretic analysis (Shannon, 1948;Nelken and Chechik, 2007) to quantify changes in stimulus encoding across drug conditions. Mutual information (MI) between single-channel MUA and vocalization stimuli was computed using two different approaches.

Using the “binwise” approach (REFs), the first 250 ms of seven vocalization stimuli were subdivided into 25 bins of 10 ms each, and each of these bins treated as one of 175 unique stimuli S. Responses R were binary (spike or no-spike) responses at the same resolution, and the MI was calculated as I(R|S) and expressed as “bits per bin.” Using the “classifier” approach (REFs), 250 ms single-trial responses were discretized into 25-letter words, with each letter determined by the spike count in a 10 ms bin. A linear discriminant classifier was used to predict one-fourth of the trials after training on the other three-fourths of trials; this process was repeated four times to generate a prediction for every trial. The predicted stimuli were treated as responses RP, and the classifier MI calculated as I(RP|S).

For both approaches, we mitigated MI bias caused by sampling limitations in the estimation of probability distributions (REFs) by using quadratic extrapolation (REF). After this procedure, we excluded from further MI analysis channels with MI values less than four times the shuffle bias or less than four times the Panzeri-Treves bias (REFs) in the control condition; we also excluded any channels that had no detectable response to any stimulus.

For statistical comparison of drug effects, we used linear mixed effects models, related to general linear models, to measure within-channel differences while accounting for the non-independence of multiple channels recorded from the same animal. We treated drug dose as a categorical fixed effect according to behavioral condition (control, sub-hypnotic, hypnotic, and recovery). We treated animals and channels as hierarchical random effects, with random slopes for dose. The baseline and driven firing rates and the mutual information were log-transformed to improve the normality of residuals. We compared models fit with drug dosage to models that omitted drug dosage using a likelihood ratio test. If this test was not significant, we concluded that there was no significant drug effect for that parameter. If there was a significant drug effect, we subsequently compared the coefficients for different dosages to determine whether subhypnotic or hypnotic doses differed from control or each other. Because residuals deviated slightly from normality for some measures, we used hierarchical bootstrapping (Brumback and Rice, 1998) of animals and channels to obtain confidence intervals that are more accurate; these confidence intervals did not differ qualitatively from those obtained under assumptions of normality, so we proceeded to assume normality when comparing coefficients. We checked residuals versus fitted values for all models and did not observe substantial heteroscedasticity.

We used a similar procedure to estimate the correlations between changes in mutual information and changes in baseline and driven rate and STTC. The outcome variable was the change in mutual information from control to hypnotic; we treated changes in the other measures from control to hypnotic as fixed effects. We also included the drug type as a fixed effect. As above, we included animals and channels as random effects, except with random slopes for drug type. We tested two types of hypotheses with this analysis. First, we tested whether there was a significant effect by drug type. If so, we could conclude that the drug had an effect on information not captured by linear relationships with the other measures. Second, we tested whether there was a significant interaction between the measures and drug type. If present, such an interaction could provide a mechanism to explain differences in information changes between drug types. We tested each type of hypothesis by comparing full models to models omitting the parameters of interest using a likelihood-ratio test.

### Histology

Electrode locations were confirmed via histological analysis, as described (Banks et al., 2011;Smith et al., 2012;Raz et al., 2014). In brief, animals were perfused intracardially with PBS and then paraformaldehyde under deep sodium pentobarbital (>90 mg/kg, IP) anesthesia, brains removed, fixed overnight, frozen and sectioned coronally into 60 µm sections. These were mounted serially on slides, stained with Cresyl Violet, and coverslipped. Electrode entry, tracks, and tip position in the brain were determined by examination of serial sections using light microscopy camera lucida techniques. Locations in the brain were identified initially using the terminology and atlas of Paxinos and Watson (Paxinos and Watson, 2007). Refer to Ref. (Smith et al., 2012) for a full description of cytological features used to aid in identification of auditory cortical areas. Electrode sites were then mapped to corresponding functionally-defined auditory areas described in Ref. (Polley et al., 2007) based on the dorsal-ventral and rostral-caudal position of the site of electrode entry.

Post-hoc histological analysis confirmed the locations of recording sites in auditory cortex (Supplementary Figure 1B-C). Of 96 electrodes across 6 animals, 81 were confined to A1 or AAF in the nomenclature of Read and colleagues (Polley et al., 2007). The electrode arrays consisted of two parallel rows (1 medial, 1 lateral) separated by 500 microns. We attempted to map each recording channel to a specific layer. In 5 of 6 animals, the medial row electrodes were located in layers 5 and 6; in four of these animals, electrodes in the lateral row were located primarily in layers 4 and 5, and in one animal primarily in layers 1 – 3. In the sixth animal, both rows of electrodes were located in layers 1 – 3.

## Results

### Anesthetic-induced loss of righting reflex

The data presented here were recorded from 6 rats implanted with microwire arrays in auditory cortex. In each recording session, data under waking conditions were recorded and then the animals were given increasing doses of anesthetic until loss of righting reflex (LORR), our surrogate measure for loss of consciousness, occurred. Mean (± SD) drug dosages for LORR were as follows: isoflurane 0.94% (±0.11); propofol 622 µg/kg/min (±50.0); dexmedetomidine 0.43 µg/kg/min (±0.050).

### Unit activity in auditory cortex

For all animals, pure-tone frequency response areas (FRAs) were recorded during the first recording session, 5 – 7 days after electrode implant (Figure 1A-C). In most cases, FRAs had clear peaks at a single best frequency (BF). In five of six animals with cortical implants, all later confirmed to have most recording sites located in A1, BFs shifted systematically from high frequencies at the rostral end of the array to low frequencies at the caudal end of the array (Figure 1D). The one exception was that in one of these five animals there was an abrupt jump in BF at the two most-caudal recording sites, likely indicating their location in PAF (Figure 1D). In the sixth animal with a cortical implant, whose recording sites were located more rostrally and ventrally in AAF, BF was constant or increased in the rostral-caudal direction. In two animals, FRAs were repeated three weeks later. Measured FRAs were largely consistent with those recorded earlier, and recorded FRAs exhibited no change in BF within the limited resolution of our stimulus set (see Methods). BFs of recorded clusters spanned the range from 4 – 64 kHz, with a concentration between 40 – 50 kHz (Figure 1E). Minimum spike latencies in response to pure tone stimuli for electrodes confirmed to be located in A1 or AAF were typically ~10 −15 msec (Figure 1F).

### Changes in firing rate across awareness states

Under control conditions, spontaneous rates of multiunit activity ranged from 0.134 to 57.4 spikes/sec (19.3 +/-13.8 spikes/sec). Consistent with previous reports (Zurita et al., 1994;Gaese and Ostwald, 2001;Hentschke et al., 2005;Hudetz et al., 2009), we observed a dramatic decrease in spontaneous spiking activity in auditory cortex with all agents, even at sub-hypnotic doses for pro and iso (Figure 2; Table 1a). Changes in spontaneous activity were most pronounced with iso, slightly less so with pro, and least with dex, but even for dex the decrease in spontaneous rate from control to just-hypnotic dose was>40% (Table 1a).

**Table 1.** Effects of anesthetic drugs on spiking properties. Values depicted are predicted means with 95% confidence intervals in brackets. For the sub-hypnotic column, stars indicate significance relative to the control condition (*p<0.05; **p<0.01; ***p<0.001) and therefore reflect drug effects rather than loss-of-consciousness. Similarly, for the hypnotic column, stars indicate significance relative to the sub-hypnotic condition, and therefore reflect dose-dependent drug effects as well as possible effects of loss-of-consciousness.

|  |  | Control | Sub-hypnotic | Hypnotic | Recovery |
| --- | --- | --- | --- | --- | --- |
| clicks | pro | 7.06 [4.79, 10.2] | 4.08 [2.92, 5.58] *** | 2.61 [1.68, 3.85] ** | 8.6 [5.67, 12.8] |
|  | iso | 8.48 [5.47, 12.9] | 4.44 [2.63, 7.17] *** | 2.42 [1.68, 3.36] *** | 10.8 [7.52, 15.3] |
|  | dex | 8.88 [5.42, 14.2] | 7.28 [3.93, 12.9] | 5.3 [2.62, 9.96] *** | 10.3 [5.91, 17.5] |

|  |  | Control | Sub-hypnotic | Hypnotic | Recovery |
| --- | --- | --- | --- | --- | --- |
| clicks | pro | 2.15 [2.06, 2.25] | 2.18 [2.06, 2.30] | 2.2 [1.89, 2.51] | 2.07 [1.86, 2.28] |
|  | iso | 2.18 [2.08, 2.28] | 2.22 [2.11, 2.33] | 2.27 [2.11, 2.42] | 2.17 [2.08, 2.26] |
|  | dex | 2.16 [2.10, 2.21] | 2.34 [2.20, 2.49] ** | 2.55 [2.35, 2.76] ** | 2.19 [2.14, 2.24] |

|  |  | Control | Sub-hypnotic | Hypnotic | Recovery |
| --- | --- | --- | --- | --- | --- |
| vocalizations | pro | 12.8 [6.58, 24.3] | 7.08 [4.00, 12.0] *** | 5.35 [2.80, 9.61] * | 12.0 [6.04, 23.1] |
|  | iso | 18.6 [12.0, 28.4] | 12.7 [8.81, 18.1] *** | 8.79 [5.77, 13.2] *** | 23.9 [15.9, 35.6] |
|  | dex | 17.5 [10.6, 28.4] | 15.7 [10.4, 23.5] | 12.4 [7.58, 19.8] *** | 19.1 [11.2, 32.0] |
| clicks | pro | 9.66 [6.12, 15.0] | 5.66 [4.12, 7.66] * | 4.08 [2.45, 6.47] * | 11.9 [7.63, 18.1] |
|  | iso | 12.2 [8.00, 18.3] | 6.58 [4.06, 10.4] *** | 4.44 [3.13, 6.18] ** | 15.8 [11.2, 22.1] |
|  | dex | 12.1 [7.16, 20.0] | 10.5 [5.88, 18.2] | 7.30 [3.38, 14.7] * | 13.7 [7.69, 23.9] |
| FM sweeps | pro | 10.9 [5.76, 19.8] | 8.07 [5.46, 11.7] | 4.60 [2.76, 7.34] *** | 9.65 [5.34, 16.9] |
|  | iso | 11.4 [4.28, 28.0] | 9.41 [4.26, 19.6] | 4.56 [1.53, 11.2] *** | 9.78 [2.74, 30.1] |
|  | dex | 10.3 [4.42, 22.6] | 8.75 [3.61, 19.6] | 6.34 [2.20, 15.8] | 9.43 [3.50, 23.2] |

|  |  | STTC |  |  |  |
| --- | --- | --- | --- | --- | --- |
|  |  | Control | Sub-hypnotic | Hypnotic | Recovery |
| vocalizations | pro | 0.129 [0.0858, 0.172] | 0.0975 [0.0628, 0.132] * | 0.0936 [0.0552, 0.132] | 0.104 [0.0703, 0.137] |
|  | iso | 0.0877 [0.0663, 0.109] | 0.093 [0.0533, 0.133] | 0.128 [0.0802, 0.176] * | 0.0902 [0.0659, 0.114] |
|  | dex | 0.081 [0.0197, 0.142] | 0.111 [0.0513, 0.17] | 0.157 [0.105, 0.209] | 0.103 [0.0766, 0.13] |
| clicks | pro | 0.0989 [0.073, 0.125] | 0.0643 [0.0452, 0.0833] | 0.0984 [0.0659, 0.131] | 0.123 [0.0942, 0.152] |
|  | iso | 0.115 [0.0815, 0.148] | 0.0695 [0.0312, 0.108] * | 0.102 [0.0665, 0.138] | 0.121 [0.093, 0.149] |
|  | dex | 0.111 [0.0796, 0.142] | 0.124 [0.0938, 0.155] | 0.146 [0.115, 0.178] | 0.142 [0.0714, 0.213] |
| FM sweeps | pro | 0.0476 [0.0418, 0.0535] | 0.0466 [0.0416, 0.0516] | 0.0541 [0.036, 0.0722] | 0.0513 [0.0405, 0.0622] |
|  | iso | 0.0559 [0.0449, 0.0669] | 0.045 [0.0344, 0.0556] * | 0.0699 [0.0366, 0.103] | 0.0716 [0.0416, 0.102] |
|  | dex | 0.0454 [0.0348, 0.0559] | 0.0618 [0.0255, 0.0982] | 0.109 [0.0784, 0.14] | 0.0369 [0.0219, 0.052] |

Response latencies to click stimuli were not significantly affected by pro or iso, but increased with dex in a dose-dependent fashion (Table 1b). However, changes in driven firing rate were substantial for most stimuli and agents (Table 1c; Figure 3). The largest effects were observed for propofol and isoflurane. Driven rates under sub doses of dex were not significantly different from those under control conditions, but there was a significant drop from sub to hyp for vocalization and click stimuli.

**Figure 3.**
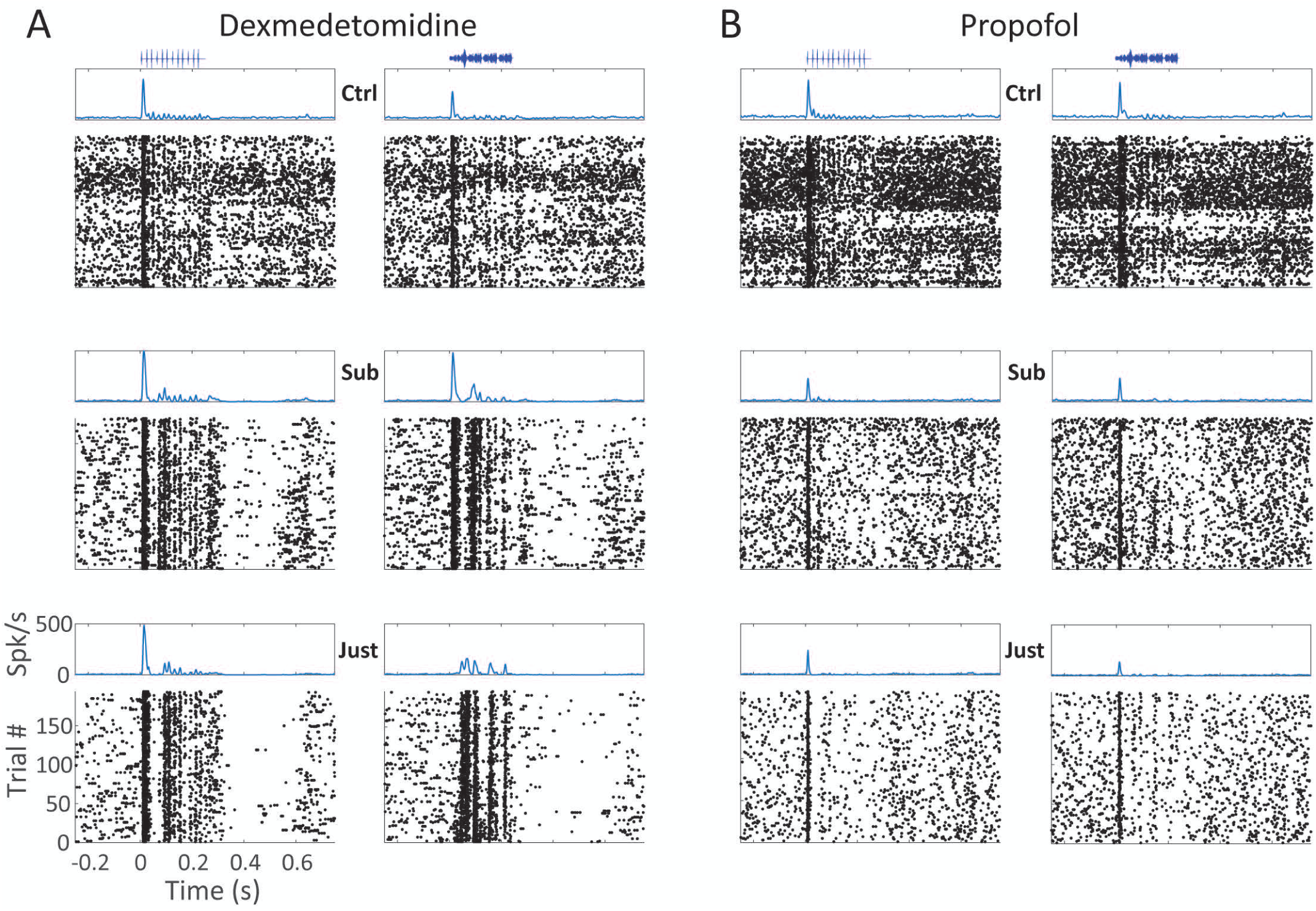
Effects of anesthetics on vocalizations and click responses. A) Instantaneous firing rate (top) and spike rasters (bottom) from one cluster showing the effect of dexmedetomidine (A) and propofol (B) on spike responses to 10 Hz click trains (left) and one vocalization stimulus (right) in control (top row), subhypnotic (middle row) and just-hypnotic (bottom row) conditions.

**Figure 3a.**
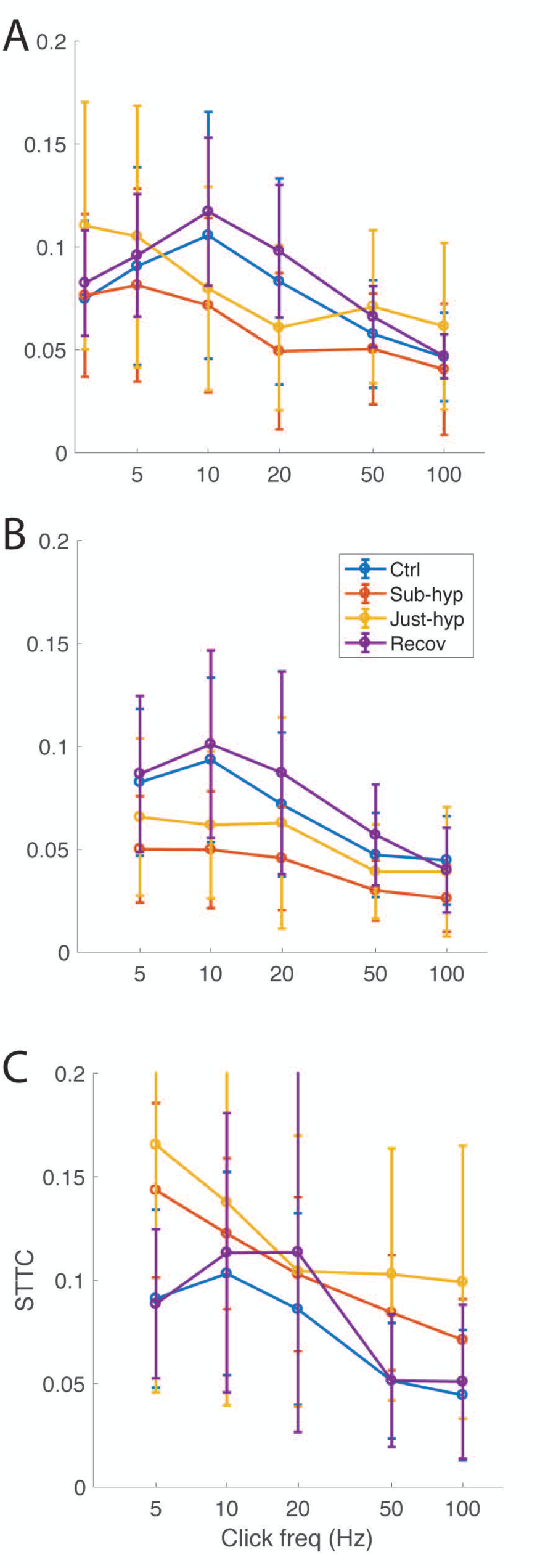
Effects of anesthetics on spike timing. Mean (+/-SD) STTC across all clusters and all animals for click trains recorded in control, sub-hypnotic, just-hypnotic and recovery for isoflurane (A), propofol (B) and dexmedetomidine (C).

### Changes in spike timing across awareness states

Cells in the ascending auditory pathway are exquisitely sensitive to temporal transients in acoustic stimuli, which carry information about their identity and location (Rosen, 1992;Huetz et al., 2011;Middlebrooks, 2015). Anesthetics at doses well above those causing LOC have several reported effects on temporal features of spiking responses, including transforming sustained into onset responses (Wang, 2007), decreasing the maximum temporal following frequency (Wang et al., 2008), and changes in mean and variability of first spike latency (Gaese and Ostwald, 2001;Ter Mikaelian et al., 2007). However, whether these effects are a common element of anesthetic agents at just-hypnotic doses has not been tested.

Unit activity in auditory cortex exhibited precise and reliable timing to specific components of all three stimulus sets (Figure 3). How spike timing varied with stimulus features and across drug conditions is mostly easily observed using click train stimuli. Clusters that were responsive to the stimuli generally followed a similar profile as click frequency was varied. At frequencies above 20 Hz, clusters were generally unable to follow click stimuli on a cycle-by-cycle basis, and onset responses became common, with some clusters also exhibiting sustained response components (not shown).

To quantify these effects, we computed the STTC (Figure 3a), the mean fraction of spikes coincident between pairs of spike trains, beyond what would be expected by chancev(Cutts and Eglen, 2014). Coincidence is measured within a specified time window, chosen here to have a relatively conservative width of 2 msec; drug effects computed using windows up to 16 msec in width were not qualitatively different (not shown). The STTC curve was typically a non-monotonic function of click frequency, with a peak at 10 Hz and decreasing at both lower and higher frequencies. Peak STTC values under control conditions averaged ~0.05 – 0.1. Effects on STTC were agent- and frequency-specific. At 5 Hz, in dex and iso (Figure 3aA, C), we observed increases in STTC at just-hypnotic doses, and for dex at the sub-hypnotic dose as well. This effect was partly due to a change in shape of the STTC versus frequency function observed under anesthesia, where the STTC curve now peaked at 5 Hz and decreased monotonically at higher frequencies. At 20 Hz, both agents caused a decrease in STTC. In propofol, we observed a decrease in STTC at all frequencies up to 50 Hz (Figure 3aB). We observed agent-specific effects on STTC of vocalization responses as well that largely paralleled the effects on low frequency click trains. Averaged over all vocalization stimuli, we observed a modest decrease in STTC for pro at both sub- and just-hypnotic doses, an increase for iso at just-hypnotic dose, and an increase for dex at both the sub- and just-hypnotic doses. The similarity of these effects to effects on low frequency click trains likely reflects the preponderance of low frequency components in the envelopes of the vocalization stimulus sets. Thus, these data indicate that anesthetics cause decreased millisecond-precision of spike timing at click frequencies >20 Hz, but that the transition from sub-hypnotic to just-hypnotic doses is not accompanied by dramatic changes in this precision. Furthermore, at low temporal frequencies in cortex, spike timing can actually be enhanced for certain anesthetic agents.

### Changes in information theoretic measures of stimulus representation across awareness states

To test directly the hypothesis that LOC is associated with degraded stimulus representation in sensory cortex, we computed the mutual information between stimuli and unit activity across awareness states. We calculated MI via two approaches: a binwise approach (Figure 4) and a classifier approach. Information rates were highly correlated between these two approaches (R^2^=0.78, p<0.0001), even across drug conditions (R^2^= 0.72, p<0.0001). Therefore, we focused the rest of our analyses only on the binwise approach. Changes in MI with anesthetic drugs were correlated across channels within individual animals.

**Figure 4.**
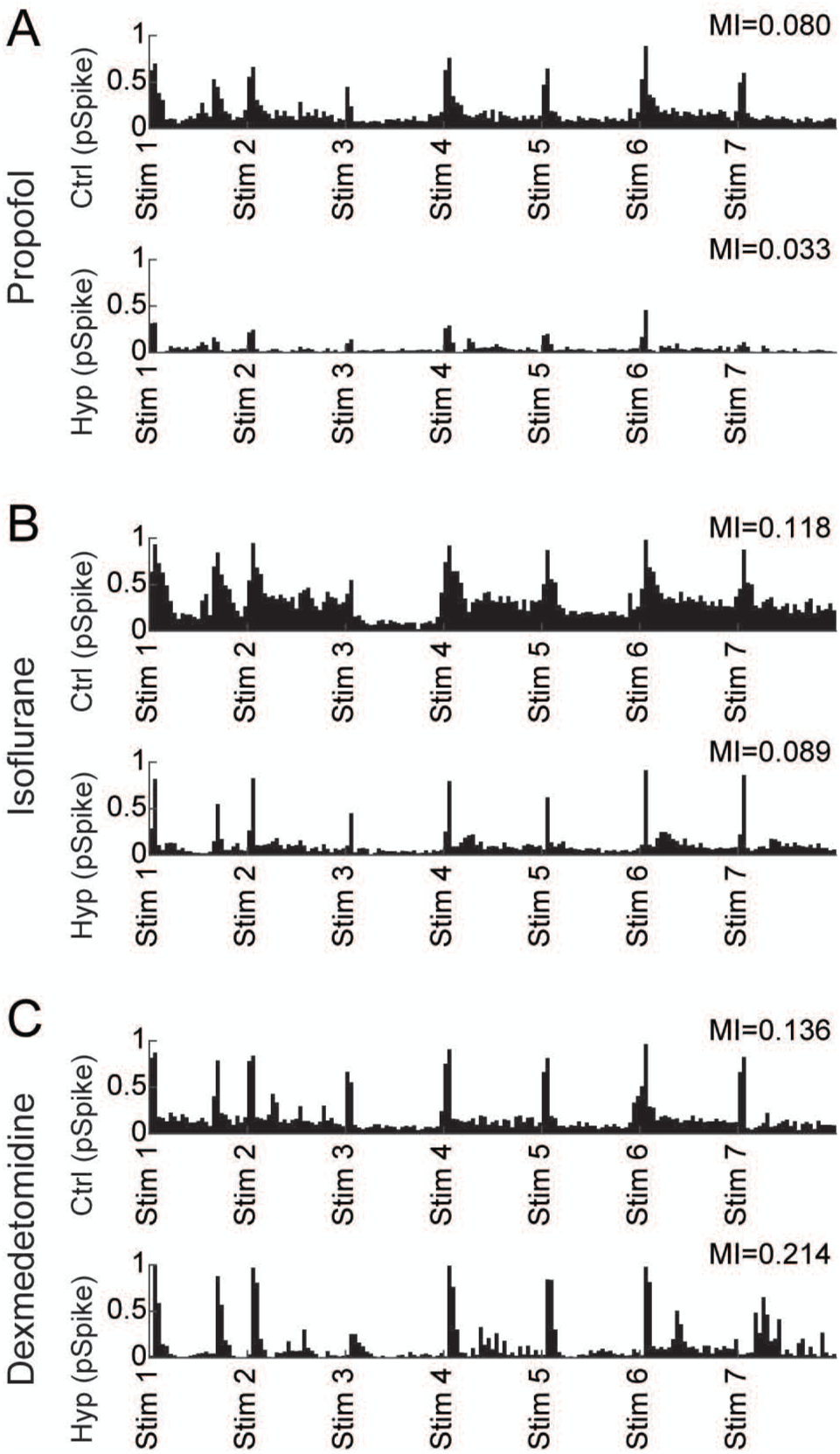
Effects of anesthetics on spike probability and mutual information. A) Probability of spiking within 10 ms bins across seven vocalization stimuli for one example channel with no drug (top) versus a hypnotic dose of propofol (bottom). MI decreased substantially in the hypnotic condition. B) Same as (A), for isoflurane. C) Same as (A), for dexmedetomidine.

We begin with the effects of anesthetics on vocalization responses (Figure 5). Two features of these data suggest that changes in stimulus information in auditory cortex are not causative for LOC. First, effects were agent-specific. MI decreased under subhypnotic doses of isoflurane and propofol, while it increased under subhypnotic doses of dexmedetomidine (Table 2). Second, if the change in brain state causing loss of consciousness was crucial to the encoding of sensory information in primary auditory cortex, we would expect moderate changes in drug concentration to effect substantial changes in MI from subhypnotic to hypnotic doses. However, for none of the agents was there a significant decrease associated specifically with LOC. For dexmedetomidine, there was in fact a significant further increase at hypnotic doses (Table 2).

**Table 2.** Effects of anesthetic drugs on binwise mutual information. Mutual information at subhypnotic levels relative to control decreased for propofol and isoflurane but increased for dexmedetomidine. Values represent model fits and 95% confidence intervals in units of bits/bin. Stars indicate significant differences relative to control for the subhypnotic column and significant differences relative to subhypnotic for the hypnotic column (*p<0.05; **p<0.01; ***p<0.001).

|  |  | Binwise Mutual Information (bits/bin) |  |  |  |
| --- | --- | --- | --- | --- | --- |
|  |  | Control | Sub-hypnotic | Hypnotic | Recovery |
| vocalizations | pro | 0.0471 [0.0339, 0.0654] | 0.0293 [0.0204, 0.0422] *** | 0.0293 [0.0195, 0.044] | 0.0372 [0.0266, 0.0519] |
|  | iso | 0.054 [0.0427, 0.0681] | 0.0411 [0.0271, 0.0622] * | 0.0378 [0.0259, 0.0551] | 0.0548 [0.0427, 0.0704] |
|  | dex | 0.0546 [0.0383, 0.0777] | 0.0781 [0.0566, 0.108] ** | 0.0965 [0.0734, 0.127] ** | 0.0683 [0.0516, 0.0904] |
| clicks | pro | 0.0447 [0.0367, 0.0546] | 0.0263 [0.0182, 0.0381] *** | 0.0324 [0.021, 0.0502] ** | 0.0522 [0.0375, 0.0728] |
|  | iso | 0.0493 [0.0375, 0.0647] | 0.0303 [0.0222, 0.0414] *** | 0.0309 [0.0205, 0.0467] | 0.0599 [0.0509, 0.0705] |
|  | dex | 0.0464 [0.0355, 0.0606] | 0.0642 [0.0543, 0.076] ** | 0.0537 [0.0284, 0.102] | 0.0479 [0.0312, 0.0733] |
| FM sweeps | pro | 0.0439 [0.0348, 0.0554] | 0.0338 [0.0224, 0.051] | 0.0335 [0.0179, 0.0625] | 0.0426 [0.0279, 0.0651] |
|  | iso | 0.0542 [0.0347, 0.0847] | 0.042 [0.0272, 0.0649] ** | 0.0469 [0.0266, 0.0828] | 0.0437 [0.0229, 0.0833] |
|  | dex | 0.0486 [0.0316, 0.0749] | 0.0566 [0.0333, 0.0963] | 0.0542 [0.029, 0.101] | 0.0342 [0.0185, 0.0632] |

**Figure 5.**
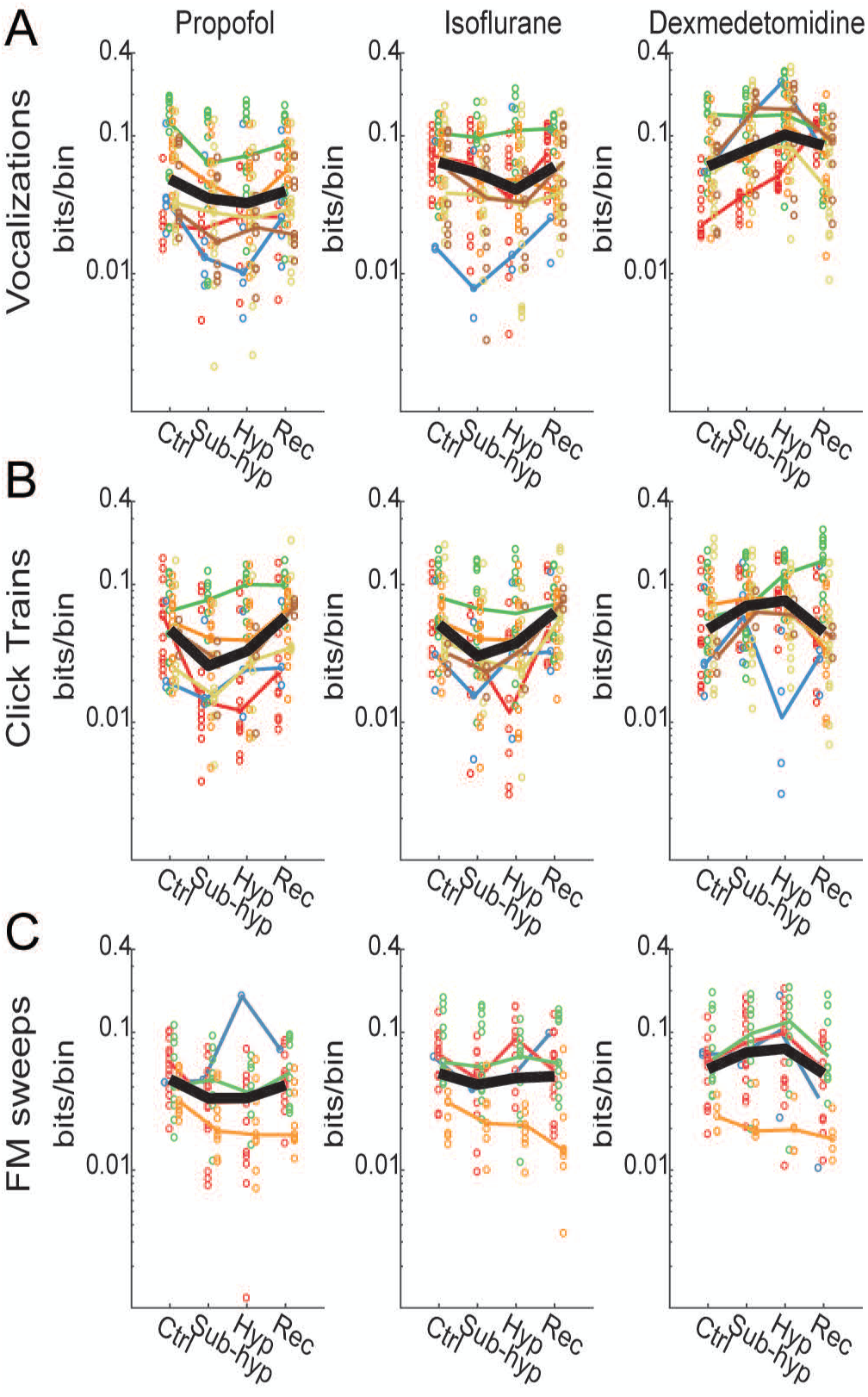
Anesthetic effects on stimulus information. Anesthetic effects were dose-dependent, but the effects of loss of consciousness varied with anesthetic agent and stimulus. A) Anesthetic effects on binwise MI for vocalization stimuli. Propofol and isoflurane caused dose-dependent decreases in MI, but dexmedetomidine showed a dose-dependent increase. B) A similar pattern was observed for click trains, except that propofol and isoflurane effects were less potent at hypnotic versus subhypnotic concentrations. C)

A similar pattern of drug effects was observed for click trains and FM sweeps, though for FM sweeps the drug effects were more modest compared to other stimuli, and only isoflurane showed a significant decrease. Thus, for all three stimuli, there decreases in MI observed under sub-hypnotic doses of propofol and isoflurane, but no further decrease observed at just hypnotic doses. For propofol, there was in a significant partial recovery of MI between the subhypnotic and just-hypnotic doses for click trains (Table 2).

For dexmedetomidine, by contrast, no decrease in MI was observed at any dose, and no consistent effect was observed across stimuli when transitioning from sub- to just-hypnotic doses. The data presented in Figure 5 suggest that changes in information are best attributed to specific drug effects rather than generalizable brain state changes occurring with loss of consciousness.

Our secondary aim was to investigate which parameters of spike trains could explain the effects on MI. Anesthetics change spontaneous and stimulus-induced spiking activity in multiple ways, including effects on firing rate, spike timing and spike variability. In control conditions, MI only correlated very weakly with baseline firing rate (Figure 6a; Table 3a). However, MI correlated somewhat with driven rate and correlated strongly with the STTC (Figure 6a; Table 3a). We also considered whether changes in baseline or driven rate or STTC would explain the changes we observed in MI. We fit linear mixed effects models (see Methods) for changes in mutual information from control to hypnotic conditions. Changes in each measure and drug type were treated as fixed effects. Interactions between measure and drug were not significant for any of the three measures (Table 3), indicating that the slope of the relationship between changes in MI and changes in baseline or driven rate or STTC were constant across drugs. We subsequently tested whether omitting the measures or the drug type affected the model fits. Both the baseline rate and STTC were significant, indicating that decreases in baseline rate and increases in STTC were correlated with increases in MI (Figure 6; Table 3). The driven rate was not significant, indicating that changes in driven rate were not associated with changes in MI. However, drug effects were significant for all three measures (Table 3), indicating that the differences in drug effects at hypnotic doses were not captured by any one of these simpler response measures.

**Table 3.**
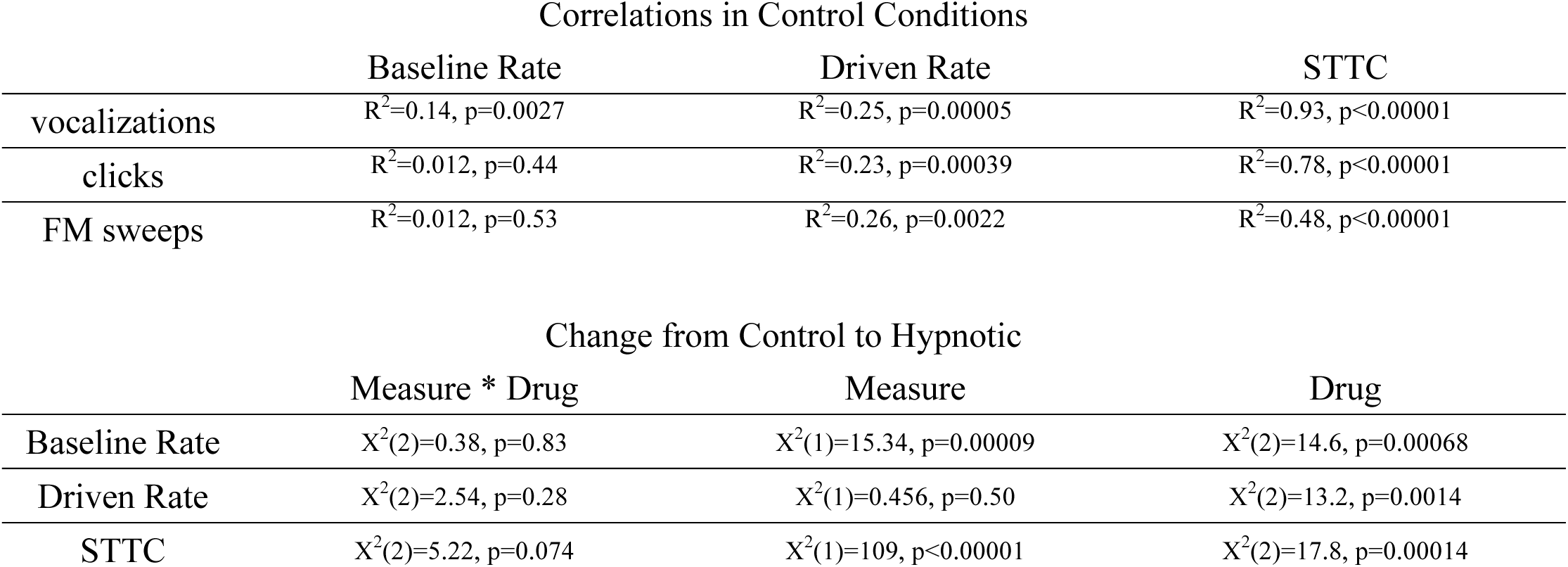
Relationship between mutual information and other measures.

**Figure 6.**
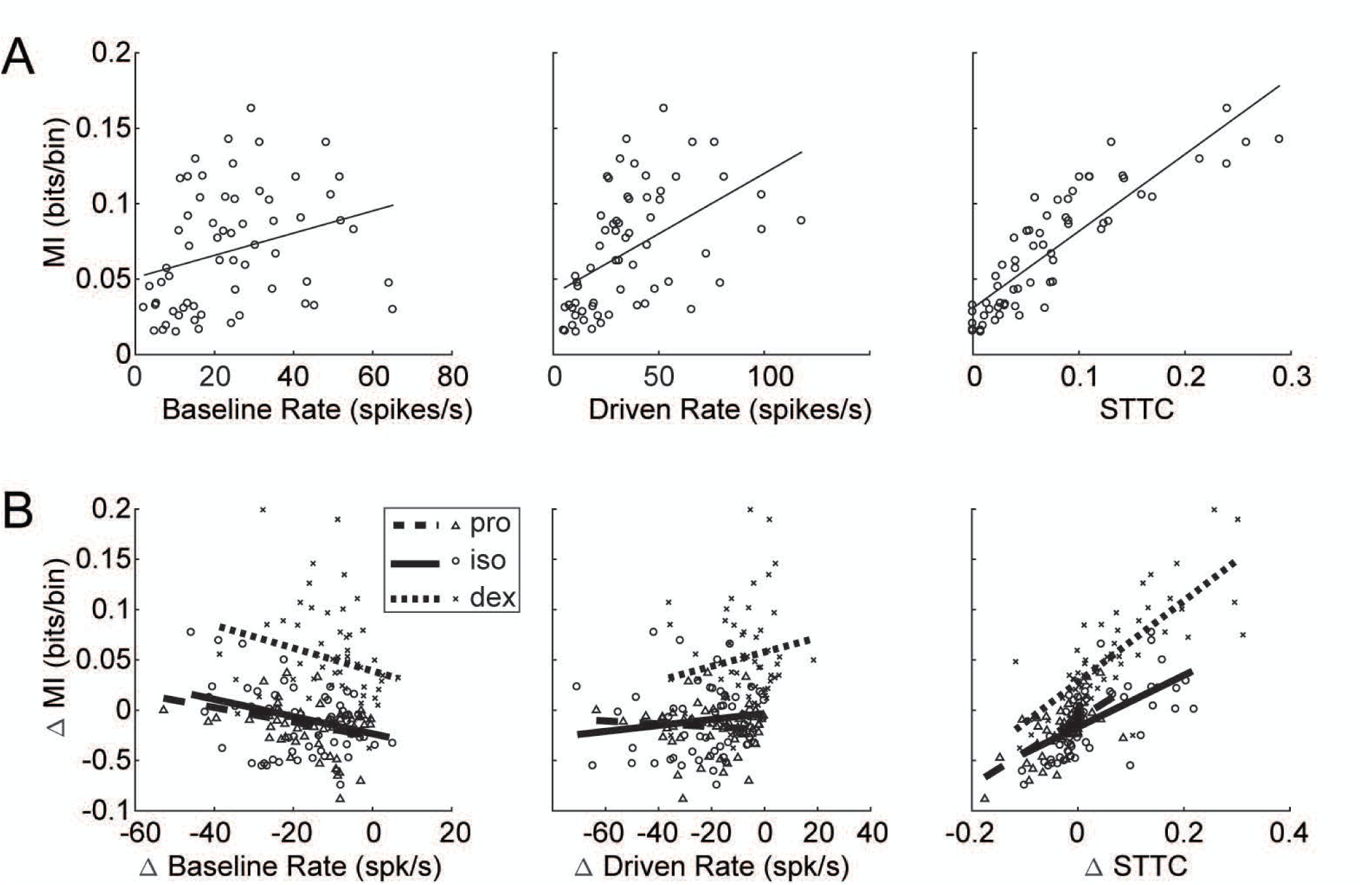
Relationship between anesthetic effect on MI and measures. A) Correlations between MI and four measures of firing with no anesthetic drug in response to vocalization stimuli. Correlations with MI were strongest for the measures that depend on spike timing: the PSTH variance and STTC. B) Relationship between the changes in firing measures and changes in MI from control to hypnotic conditions. Lines are fits from linear mixed effects models (see Methods and Table 3). Slopes did not vary significantly across drug types, but intercepts did. (other stimuli in a table).

## Discussion

### Summary

We investigated two simple questions concerning the causal role of changes in cortical sensory responses in anesthesia LOC: do diverse anesthetic agents cause consistent degradation in stimulus representation, and are the effects at just-hypnotic doses dramatically different from effects at sub-hypnotic doses? The data presented here clearly indicate that the answers to both questions is no. We observed divergent effects of three anesthetic agents on the MI between stimulus and spiking responses in auditory cortex for multiple classes of acoustic stimuli, with one agent, dexmedetomidine, increasing the information content at sub-hypnotic and just-hypnotic doses, and two others, propofol and isoflurane, decreasing the MI. In addition, effects at sub-hypnotic and just-hypnotic doses were not consistently distinct. This diversity was paralleled by diverse effects on a broad array of metrics of neural activity, and for the one metric for which we observed common effects, i.e. spontaneous firing rate, there was no dramatic difference between effects of sub-hypnotic and just-hypnotic doses. Likely because we were testing lower doses of anesthetics than in previous studies, we observed some effects that were qualitatively different than those observed in previous reports, including enhanced spike timing precision under dex and iso for some stimuli.

### Ascertaining levels of consciousness

One of the challenges of studying the mechanisms of loss and recovery of consciousness under anesthesia, and the neural correlates of consciousness in general, is ascertaining whether the subject is unconscious, as opposed to disconnected from the environment or simply unmotivated to respond. This problem is especially acute in animal studies, in which we need to rely on surrogates for LOC such as the absence of righting reflex. LORR is widely used as an indicator of LOC in rodent studies, based primarily on the remarkably strong correlation across agents between doses required to cause LOC in humans (assayed by absence of response to verbal command) and those that cause LORR in rodents (Franks, 2008). We emphasize that rodent studies such as ours, the equivalence across agents in the behavioral endpoint of LORR does not necessarily correspond to an equivalence in state of consciousness. However, the main result of this study, that impairment of stimulus representation in sensory cortex is not predictive of LOC under anesthesia, is likely to hold even the absence of a strict equivalence along this dimension. We observed *increases* in MI under dexmedetomidine compared to control at both doses tested, and for none of the agents was there a *decrease* in MI at the higher doses tested, while in all cases animals were clearly less responsive to their environment at these higher dose compared to the lower dose or wake. In the absence of any indication that higher doses of anesthesia are accompanied by suppression of stimulus information, even if LORR is not strictly associated with LOC in all cases, there is still no support for the hypothesis of a causal link between impaired stimulus representation and LOC under anesthesia.

We also note that dexmedetomidine is an atypical anesthetic. It is thought to induce a state that is similar to natural sleep (Noreika et al., 2011;Baker et al., 2014), based on its actions in the subcortical arousal centers (Nelson et al., 2003;Zhang et al., 2015), and its molecular mechanism of action. In patients, sedation with dexmedetomidine is routinely associated with LOC, as assayed by failure to respond to verbal command, but patients are still rousable (Venn and Grounds, 2001;Sanders and Maze, 2012). In addition, the incidence of dreaming, a form of conscious experience characterized by disconnection from the environment, is reported to be quite high under dexmedetomidine (Noreika et al., 2011;Sanders et al., 2012). In our experiments, rats experienced LORR under dexmedetomidine as under propofol and isoflurane, though in some instances testing for LORR could produce temporary arousal in the form of brief (<1 min) periods of directed movement, but never recovery of righting. Left undisturbed, no voluntary movement was observed.

### Changes in unit activity do not predict awareness states

Decreased spontaneous activity in cortex is a common finding across anesthetic agents (Zurita et al., 1994;Gaese and Ostwald, 2001;Hentschke et al., 2005;Hudetz et al., 2009). These observations are manifestations of suppressive effects on ongoing cortical network activity common under anesthesia, consistent with previous reports of anesthetic effects on cortical connectivity [REFs]. Suppressed network activity is likely to have an impact on stimulus representation as well; indeed, if spontaneous activity is considered strictly as ‘noise’ during the encoding of the ‘signals’ evoked by experimental stimuli, one would expect that suppressed spontaneous activity would work to enhance stimulus representation. That this expectation was not met for isoflurane and propofol does not necessarily argue against this line of reasoning; it could be that effects on stimulus-evoked activity simply overwhelmed whatever gain in MI might emerge from suppression of background spiking. In either case, the observation that substantial changes in spontaneous rate are observed even at sub-hypnotic doses is consistent with the idea that these effects on network activity and stimulus representation are not causal for LOC.

Reported effects of anesthetics on cortical sensory responses are mixed. In some studies, components of evoked responses were suppressed under anesthesia (Ikeda and Wright, 1974, Villeneuve, 2003 #7897, Hudetz, 2009 #7130), whereas other studies found substantial enhancement of evoked responses (Banks, 2010, Imas, 2005 #6155). [Also Guo et al 2012; Effects on unit activity: Sellers 2015 JNeurophys (absence of effect on spont in V1; spike timing effects). Durand et al 2016 no effect on spont, evoked rate. Hudetz. Bartlett & Wang.]

One observation of note in the present study is preserved responsiveness to acoustic stimuli under dexmedetomidine (Figure XX). By contrast, a previous study using fMRI and PET imaging in human volunteers (Akeju et al., 2014) found that LOC under dexmedetomidine was associated primarily with suppressed thalamo-cortical, rather than cortico-cortical, connectivity, consistent with the thalamic switch hypothesis (Alkire et al., 2000). Our results are consistent with dexmedetomidine inducing a sleep-like state, as evidenced by its actions on subcortical sleep and arousal centers (Nelson et al., 2003;Zhang et al., 2015) and the similarity between electrophysiological changes associated with sleep and dexmedetomidine (Baker et al., 2014).

Under reliably hypnotic doses, short latency responses to best frequency tones are comparable to awake conditions, but frequency tuning is sharper and receptive fields less complex (i.e. more like subcortical tuning) (Abeles and Goldstein, 1972, Brugge, 1985 #4519, deCharms, 1998 #4618, Gaese, 2001 #4860, Sally, 1988 #7001, Zurita, 1994 #5818). Cells in anesthetized animals have reduced ability to follow rapid temporal transients (Goldstein et al., 1959, Wang, 2008 #7218) and tend to produce onset responses, with reduced sustained components to prolonged stimuli (Gaese and Ostwald, 2001, Ter Mikaelian, 2007 #6661, Wang, 2005 #6284). Sensory responses are more stereotyped across trials and across stimuli (Gaese and Ostwald, 2001, Kisley, 1999 #4942, Ter Mikaelian, 2007), perhaps secondary to suppression of spontaneous activity by anesthetic agents and thus reduced intrinsic network “noise”. [White et al 2012 found higher variability, as did Gur 1997. Kisley & Gerstein 1999 found non-monotonic chances across anesthetic levels (highest at medium), but did not compare to awake state.]

[Effects on latency: Gaese & Ostwald 2001 found increased latency. Ter-Mikelian et al 2007 found large (12 msec) decrease in first spike latency under anesthesia, also reduced SD of first spike, i.e. onsets more precise in anesthesia. Strauss et al 2015 PNAS show that latencies of ERPs in auditory cortex increase during sleep. Relate this to less efficient info transmission from thalamus when operating in burst mode, and thus slower evidence accumulation and longer spike latencies. Cite Friston paper for theory, and two other ERP papers for confirmation of this result. See their Refs 42-44.]

### Stimulus representation during anesthesia LOC

As noted above, there have been suggestions that dexmedetomidine has effects on consciousness distinct from other anesthetic agents. For example, it was theorized recently that dexmedetomidine impairs arousal level, rather than content of consciousness (Noreika et al., 2011;Mashour and Hudetz, 2017). Evidence in support of this idea arises from a study showing that thalamo-cortical connectivity is impaired under dexmedetomidine (Akeju et al., 2014), in contrast to most reports on volatile agents and propofol (Lee et al., 2013b) [REFs]. However, our results showing the absence of suppressive effects on stimulus representation under dexmedetomidine is inconsistent with this idea. That is, impaired thalamo-cortical connectivity would be expected to reduce neural responsiveness to sensory stimuli, and to impair information content of spike trains evoked by those stimuli. We find neither to be the case.

Both methods employed to measure MI are based primarily on "rate modulation", that is, the number and reliability of the spikes as well as the ability to distinguish those numbers from those occurring at other times. Spike timing clearly enters into this analysis, but precise spike timing, as measured by STTC, or as employed in previous MI analyses of responses in auditory cortex (Chechik et al., 2006). Although changes in firing rate are known to be critical to information measures, we found that the response measures most predictive of information changes involved the temporal dynamic range and temporal reliability of stimulus responses.

Our observations are consistent with several imaging studies investigating responses to acoustic stimuli in human volunteers. For example, fMRI BOLD responses to spoken words in primary auditory cortex are suppressed under propofol sedation, i.e. doses at which response to verbal command is intact, but further suppression of these responses at slightly higher doses of propofol causing LOC are quite modest (Plourde et al., 2006). In addition, our observations are consistent with reports that changes in sensory responses under moderate doses of anesthesia are modest (Guo et al., 2012;Durand et al., 2016), and that the largest changes in sensory responses under anesthesia occur in higher order cortical areas (Liu et al., 2012). Studies in coma patients indicate that activation of primary sensory cortex by sensory stimuli persists even in the vegetative state, suggesting that activation of primary sensory cortex is necessary, but not sufficient, for consciousness (Crick and Koch, 1995;Laureys et al., 2000;Laureys et al., 2002;Boly et al., 2004). These reports, along with our observations that LOC is not associated with degraded stimulus representation in sensory cortex, all support the model in which sensory information activates primary sensory cortex, but fails to be incorporated into the cortical sensory hierarchy and enter into awareness.

### Functional implications

Responses of individual cells are likely only modestly affected by doses of anesthetics causing LOC, and these responses do not change dramatically at the transition between awake-sedation (sub-hypnotic) and hypnosis (just-hypnotic). Clearly, however, anesthesia involves sensory disconnection and the absence of sensory awareness, and the locus of mechanisms underlying these effects are likely in the cortico-thalamic network. It is possible that while population coding of sensory stimuli does not exhibit common effects across agents at anesthesia LOC, population coding may still exhibit these effects, for example due to increased redundancy across the cell assemblies encoding specific stimuli or suppressed feedback modulation of sensory responses, both possibly secondary to changes in local or long range cortical connectivity. It is also possible that the relevant locus for these common effects resides in a higher order cortical area, and that anesthesia LOC triggers roadblocks that present stimulus representations in primary cortex from entering into the cortical hierarchy. Future experiments and analyses aimed at addressing these questions will likely shed light on this important question.

## Acknowledgements

Supported by National Institutes of Health (R01 GM109086 to M. I. Banks), and the Department of Anesthesiology, School of Medicine and Public Health, University of Wisconsin, Madison, WI.

## Figure Captions

**Supplementary Figure 1.**
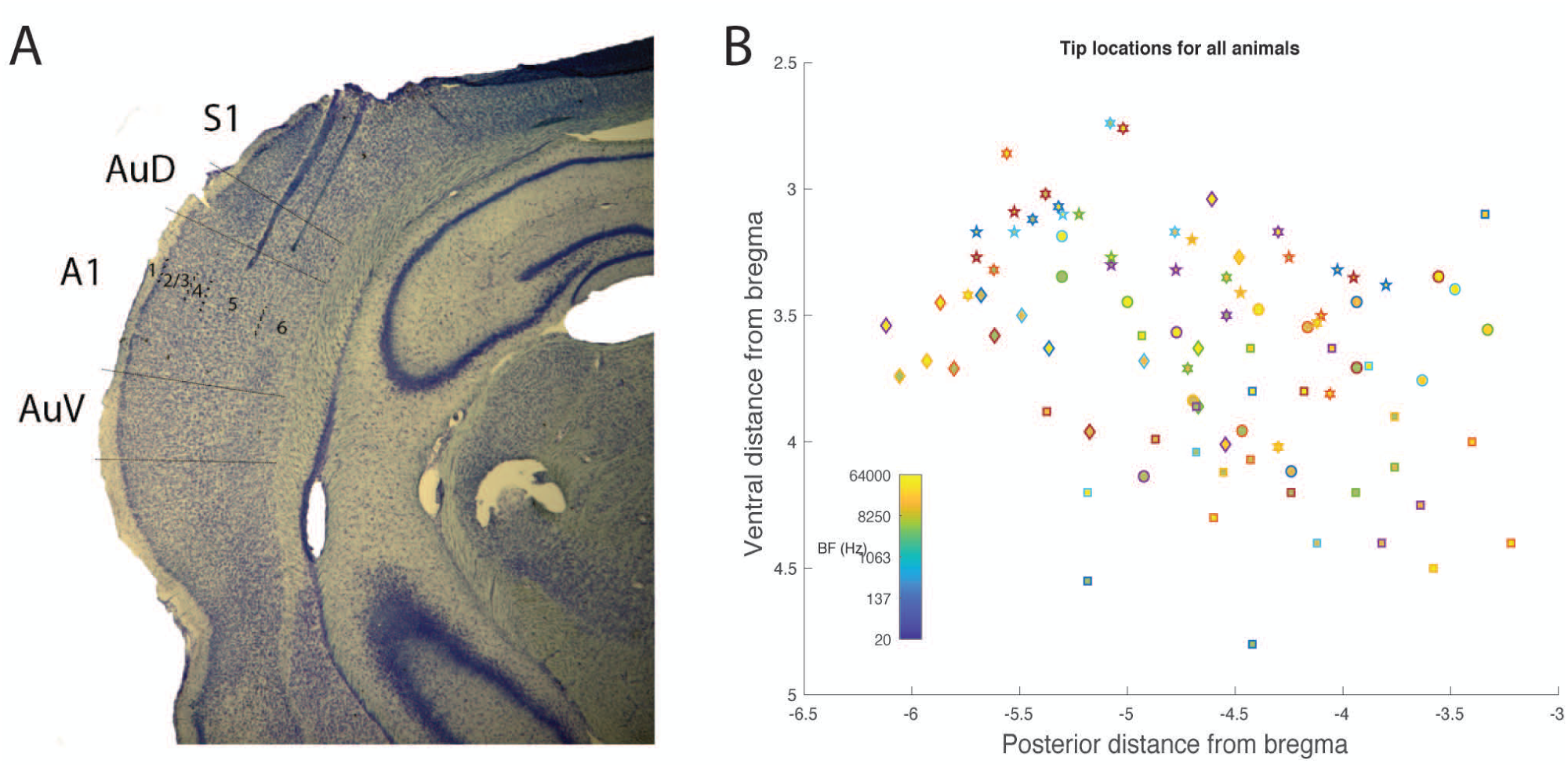
Histological analysis. A) Example photomicrograph from one animal showing two electrode tracks, with tips in infragranular layers of auditory cortex. B) Tip locations relative to Bregma for all six animals. Each symbol is a different animal. BF is indicated by color.

**Supplementary Figure 2.**
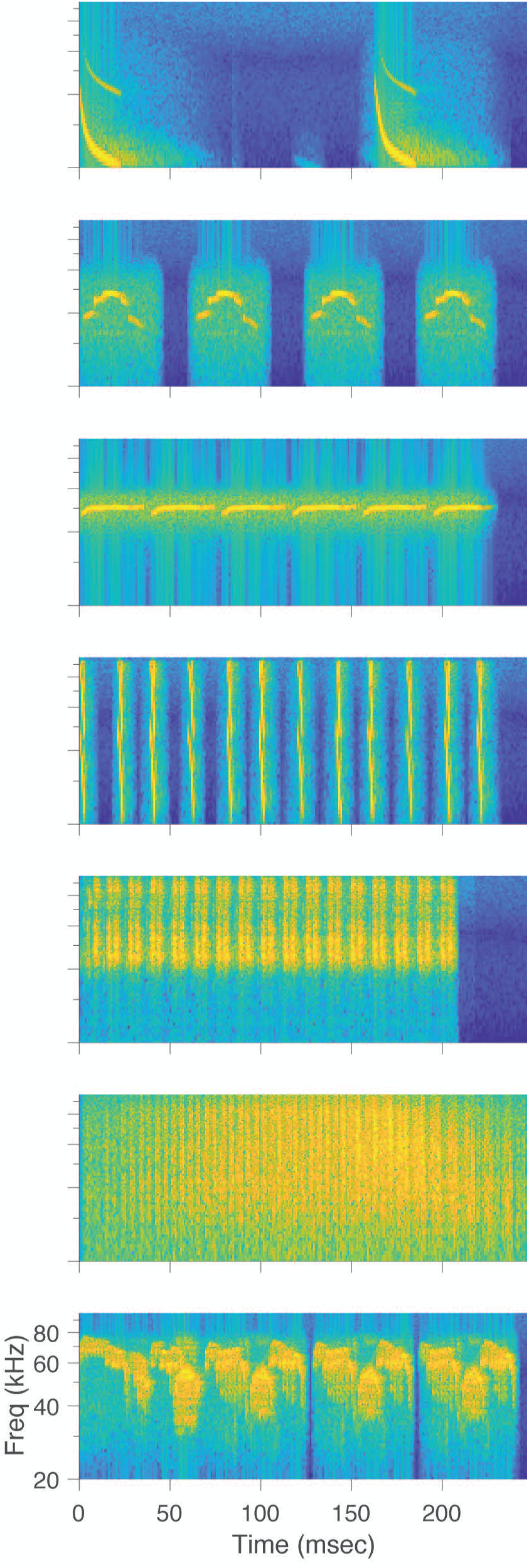
Spectrograms of vocalization stimuli.

